# Natural killer cell immunosuppressive function requires CXCR3-dependent redistribution within lymphoid tissues

**DOI:** 10.1101/2021.05.11.443590

**Authors:** Ayad Ali, Laura M. Canaday, H. Alex Feldman, Hilal Cevik, Michael T. Moran, Sanjeeth Rajaram, Nora Lakes, Jasmine A. Tuazon, Harsha Seelamneni, Durga Krishnamurthy, Eryn Blass, Dan H. Barouch, Stephen N. Waggoner

## Abstract

Natural killer (NK) cell suppression of T cells is a key determinant of viral pathogenesis and vaccine efficacy. This process involves perforin-dependent elimination of activated CD4 T cells during the first three days of infection. Although this mechanism requires cell-cell contact, NK cells and T cells typically reside in different compartments of lymphoid tissues at steady state. Here, we show that NK-cell suppression of T cells is associated with a transient accumulation of NK cells within T cell-rich sites of the spleen during lymphocytic choriomeningitis virus infection. The chemokine receptor CXCR3 is required for relocation to T-cell zones and suppression of antiviral T cells. Accordingly, this NK-cell migration is mediated by type I interferon (IFN)-dependent promotion of CXCR3 ligand expression. In contrast, adenoviral vectors that weakly induce type I IFN and do not stimulate NK-cell inhibition of T cells also do not promote measurable redistribution of NK cells to T-cell zones. Provision of supplemental IFN could rescue NK-cell migration during adenoviral vector immunization. Thus, type I IFN and CXCR3 are critical for properly positioning NK cells to constrain antiviral T-cell responses. Development of strategies to curtail migration of NK cells between lymphoid compartments may enhance vaccine-elicited immune responses.

## Introduction

Natural killer (NK) cells are innate lymphocytes with a critical role in immune defense against viral pathogens in both mice and humans (1-3). In addition to killing virus-infected cells, NK cells control the magnitude and quality of antiviral T- and B-cell responses via perforin-dependent elimination of activated CD4 T cells during the first few days after infection (4, 5). Lymphocytic choriomeningitis virus (LCMV) is a potent trigger of the immunoregulatory functions of NK cells with significant consequences regarding viral clearance and development of immune pathology (4-6). Similar mechanisms constrain humoral and cellular immunity after immunization of mice (7, 8). NK-cell suppression of T-cell or antibody responses is also apparent in humans during infections with HIV or hepatitis B virus (HBV) and after administration of yellow fever, malaria, or HBV vaccines (9-13). An improved understanding of the mechanisms essential for NK-cell suppression of adaptive immune responses is likely to facilitate development of interventions to enhance the efficacy of vaccines or amplify immune responses during virus infection.

NK-cell suppression of T cells has been reproducibly linked to perforin (4-6, 8, 14), a key component of a cell-contact dependent granule-mediated killing (15). Thus, the immunoregulatory activity of NK cells likely involves physical liaisons with T cells in lymphoid tissues. Yet in the absence of inflammation, NK cells are infrequently present at T-cell rich sites, including lymph nodes and the white pulp of the spleen (16-20). Moreover, NK cells present in human lymph nodes typically express little perforin (21). However, infections with murine cytomegalovirus (MCMV) or LCMV trigger NK-cell accumulation in the white pulp of spleen (16-18), while infections of humans and non-human primates with HIV or SIV result in significant localization of NK cells into T/B-rich lymphoid follicles (22). In this study, we establish that positioning of NK cells within T-cell rich white pulp follicles is essential for NK-cell mediated suppression of antiviral T-cell responses.

## Results

### Transient positioning of NK cells in T-cell zones during infection

NK cells eliminate a fraction of virus-specific CD4 T cells in a perforin-mediated, putatively cell-contact-dependent manner in the spleen during the initial three days of LCMV infection in mice (4, 5). However, NKp46^+^ NK cells predominately populate the red pulp regions of the mouse spleen with few NK cells detectable in the T-cell rich white pulp at baseline (**Figure 1A, Figure S1**). Consistent with previous reports (16, 18), infection with LCMV triggered increased proportions (**Figure 1B-C**) and numbers (**Figure 1D**) of NKp46^+^ NK-cell localizing within the splenic white pulp (**Figure 1B, D**) and T-cell zones (**Figure 1C**). This re-localization was apparent as early as 24 hours and continued to increase until roughly 60% of splenic NK cells localized within the white pulp by day 3 of infection (**Figure 1A-D**).

**Figure 1.**
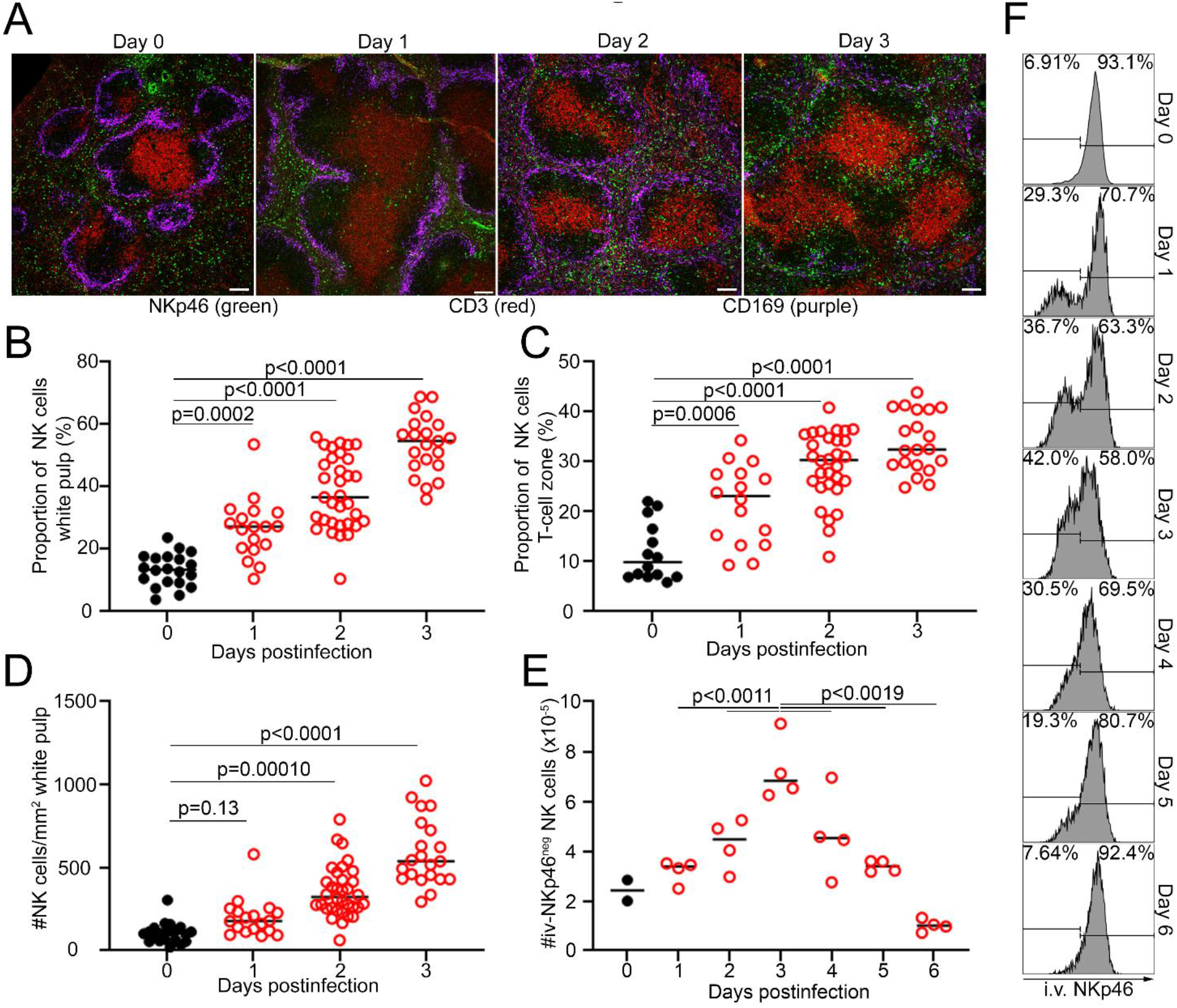
Transient positioning of NK cells in T-cell zones during infection. (A-F) C57BL/6 mice (n=3-4/group) were infected with the Armstrong strain of LCMV. The proportion (B, C) and number (D) (5-12 follicles/mouse) of NKp46^+^ NK cells (green) enumerated within (B, D) CD169 macrophage (purple) bound white pulp or in (C) CD3^+^ T-cell zones (red) is plotted. (E, F) At indicated times after infection, mice were intravenously injected with anti-NKp46 antibody 3 minutes prior to euthanasia to label splenic NK cells (CD3^neg^TCRβ^neg^CD8α^neg^CD49b^+^NK1.1^+^ex-vivoNKp46^+^) in vascularized red pulp (iv-NKp46^+^) or white pulp (iv-NKp46^neg^) regions. Mean (E) number of iv-NKp46^neg^ (white pulp) NK cells is graphed at each time point (4 mice/time) along with (F) representative histograms of iv-NKp46 staining of gated NK cells. Data reflect one of two independent experiments with statistical differences determined by one-way ANOVA. Scale bars measure 100 µm.

To more precisely quantify NK-cell localization, we took advantage of differences in vascularity between spleen red pulp and white pulp that can be detected using a modification of established intravascular staining methods (23). We intravenously injected an APC-labeled anti-NKp46 antibody and euthanized mice after 3 minutes. Ex vivo staining with anti-NKp46 ubiquitously labels splenic red pulp NK cells (**Figure S2A-B**) in both infected and uninfected mice (24). Consistent with restricted labeling of NK cells in poorly vascularized sites following intravenous injection of antibody (iv-NKp46), NK cells in the white pulp were shielded from intravascular staining but readily labeled with anti-NKp46 antibodies applied to tissues sections ex vivo (**Figure S2C**). Thus, both microscopy (**Figure 1A**) and intravascular staining (iv-NKp46^neg^, **Figure 1E, F**) revealed few white pulp localized NK cells and a predominance of red pulp localized NK cells (**Figure 1F, Figure S1**) in the absence of virus. Following LCMV infection, the fraction (**Figure 1F**) and number (**Figure 1E**) of splenic iv-NKp46^neg^ NK cells increased over time, peaking at day 3 and returning to near baseline by day 6 of infection. Notably, differences in iv-NKp46 staining could not be explained by differences in NKp46 expression levels (ex vivo stain, **Figure S2A**). In total, these results show that NK-cells transiently re-locate to the splenic white pulp during the first three days of LCMV infection, a window of time concomitant with perforin-dependent NK-cell killing of activated CD4 T cells (5).

The intravascular staining method permitted comparison of the phenotype of NK cells in the white pulp (iv-NKp46^neg^) and red pulp (iv-NKp46^+^) after infection. Each subset exhibited similar expression levels of activating (Ly49H, DNAM-1, NKG2D) and inhibitory (CD94, NKG2A, KLRG1) NK-cell receptors, although NKG2D expression was slightly reduced on white pulp NK cells (**Figure S3A-F**). Expression of CXCR3 (**Figure S3G**) as well as the distribution of immature (CD11b^neg^ CD27^+^), transitional (CD11b^+^ CD27^+^), and mature (CD11b^+^ CD27^neg^) NK cell subsets (**Figure S3H**) were also similar between the red and white pulp. However, white pulp NK cells expressed higher levels of granzyme B and the IL-2 receptor alpha (CD25) than red pulp NK cells (**Figure S3I-J**). Thus, white pulp NK cells may be more cytolytically active and IL-2 responsive than their counterparts in the red pulp.

### Type 1 interferons are necessary and sufficient to drive NK cell localization in white pulp

In contrast to LCMV infection, vaccination with a replication incompetent adenovirus serotype 5 vector (Ad5) triggered robust T-cell responses without any evidence of T-cell suppression by NK cells (25, 26). We hypothesized that the absence of NK-cell immunoregulatory functions after Ad5 vaccination may be associated with weak or absent induction of NK-cell localization within splenic T-cell zones. As such, we immunized mice with replication incompetent Ad5 vectors harboring either the glycoprotein (GP) of LCMV (25) (**Figure S4**) or β-galactosidase (Ad5-LacZ) (**Figure 2**) and assessed NK-cell follicular localization. There was no measurable localization of NK cells within the white pulp at any time point measured after Ad5 immunization (**Figure 2A-B, Figure S4A-B**), supporting an association between NK cell migration and immune regulation.

**Figure 2.**
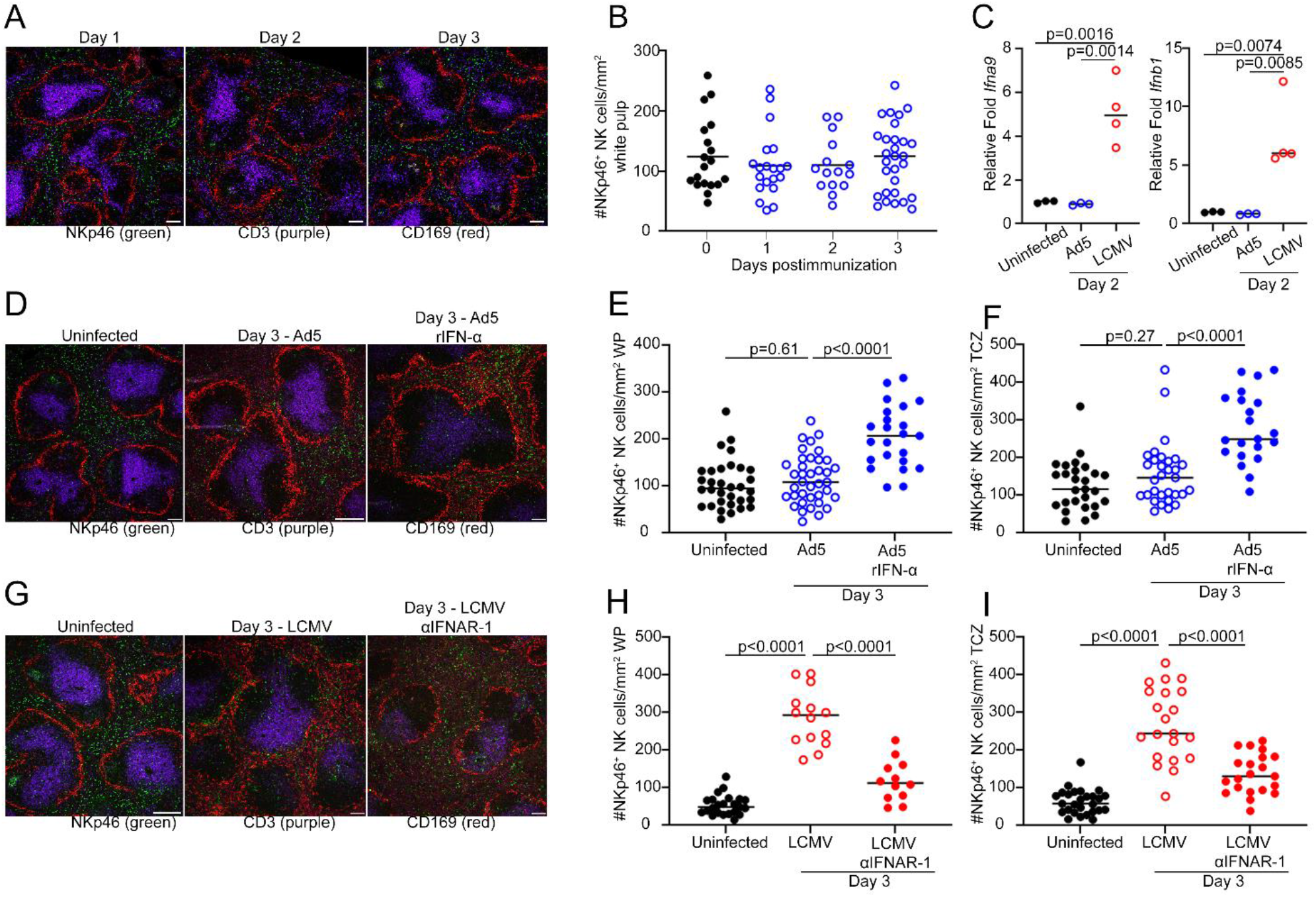
Type 1 interferons are necessary and sufficient to drive NK cell white pulp localization. C57BL/6 mice were inoculated with (A-B) Ad5-GP, (C-F) Ad5-LacZ, or (C, G-I) LCMV. One group (D-F) of Ad5-LacZ inoculated C57BL/6 mice were treated with 8 µg/day recombinant IFN-α while another group (G-I) of LCMV-infected C57BL/6 mice were treated with 400 µg/day anti-IFNAR-1 antibodies. At indicated times post inoculation, spleens were imaged (representative images shown in A, D, G) to determine localization of NKp46^+^ NK cells (green) relative to CD3^+^ T cells (purple) and CD169^+^ macrophages (red), with enumeration of NK cells in (B, E, H) CD169-delineated white pulp or (F, I) CD3^+^ T-cell zone in 5-12 follicles per mouse (n=3-4 mice/time point). (C) Relative expression of *Ifna9* and *Ifnb1* two days following LCMV infection or Ad5-LacZ inoculation compared to uninfected mice. Data reflect one of two independent experiments with statistically significant differences determined by one-way ANOVA. Scale bars measure 100 µm.

One important distinction between LCMV infection and Ad5-vector immunization lies in the magnitude of type I interferon (IFN-I) responses, where LCMV is a potent inducer of IFN-I (27). Indeed, we measured a ∼5-fold increase in *Ifna9* and *Infb1* expression during LCMV infection but not Ad5 immunization (**Figure 2C**). Thus, we hypothesized that robust IFN-I expression during LCMV infection but not after Ad5 immunization promotes NK cell follicular migration. Indeed, daily provision of recombinant IFN-alpha (rIFN-α) during Ad5 immunization resulted in increased NK-cell localization within the white pulp (**Figure 2D-E**) and T-cell zones (**Figure 2D-F**) of the spleen, as well as enhanced NK-cell accumulation in the draining lymph nodes (**Figure S5**). In agreement with the importance of IFN-I for NK-cell migration, addition of anti-IFNAR-1 blocking antibodies during LCMV infection impeded localization of NK cells in the white pulp (**Figure 2G-H**) and T-cell zones (**Figure 2G-I**) of the spleen. IFN-I blockade also reduced NK-cell accumulation in draining lymph nodes after LCMV infection (**Figure S5**). Thus, IFN-Is are important triggers of NK cell migration to T-cell rich sites in lymphoid tissues.

### NK cells require CXCR3 for splenic T-cell zone localization

Inflammatory recruitment of NK cells to lymph nodes during poxvirus infection or after dendritic cell immunization requires expression of the chemokine receptors CXCR3 on NK cells (20, 28, 29). Likewise, CXCR3 is implicated in NK-cell localization to the white pulp after polyinosinic:polycytidylic acid (poly I:C) injection or MCMV infection (16, 17). Therefore, we hypothesized that type I IFN induction of CXCR3 ligands and CXCR3 expression on NK cells is vital for migration to T-cell zones in the splenic white pulp. The expression of the CXCR3 ligands *Cxcl9, Cxcl10* and *Cxcl11* were elevated following LCMV infection but not Ad5 immunization (**Figure 3A**). IFNAR-1 blockade dampened LCMV-mediated induction of *Cxcl10* and *Cxcl11* (**Figure S6**).

**Figure 3.**
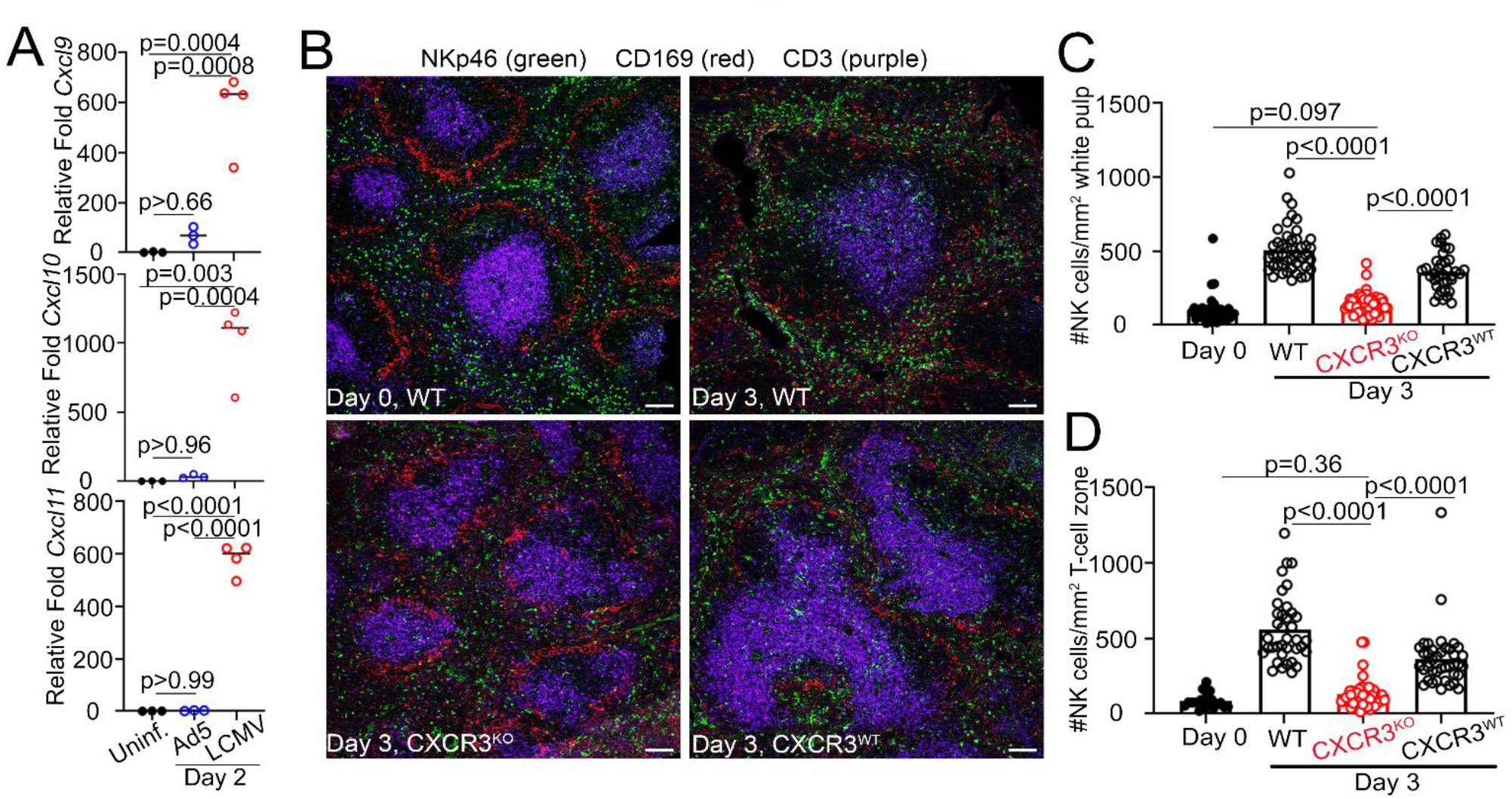
NK cells require CXCR3 for splenic T-cell zones localization. (A) Relative expression of *Cxcl9, Cxcl10 and Cxcl11* in the spleens of uninfected, Ad5-LacZ inoculated or LCMV-infected C57BL/6 mice (n=4/group). (B-D) C57BL/6 mice (WT) and mixed CXCR3^WT^ or CXCR3^KO^ bone-marrow chimeras (n=3) were infected with LCMV. Prior to (Day 0) or after (Day 3) infection, (B) confocal microscopy was used to determine median number (5-12 follicles/mouse) of NKp46^+^ NK cells (green) in (C) CD169^+^ macrophage (red) bordered white pulp or in (D) CD3^+^ T-cell zone (purple). Data reflect one of two independent experiments with statistically significant differences determined by one-way ANOVA. Scale bars = 100 µm.

To address the role of CXCR3 in NK cell migration and suppressive function without undermining the role of CXCR3 in T-cell activation (30), we generated mixed bone marrow chimeric (BMC) mice harboring CXCR3-sufficient T and B cells in conjunction with an innate compartment that is either CXCR3-sufficient or –deficient. *Rag*^*KO*^*Cxcr3*^*WT*^ or *Rag*^*KO*^*Cxcr3*^*KO*^ bone marrow cells (Ly5.2) were mixed at a 9:1 ratio with wild-type bone marrow cells (Ly5.1) prior to reconstitution of lethally irradiated mice. This protocol (31) ensures RAG-dependent cells, including T and B cells, are ubiquitously CXCR3-sufficient while innate cells like NK cells are predominately derived from the *Rag*^*KO*^ precursors that are either *Cxcr3-*sufficient or - deficient. We confirmed that CXCR3 was absent from NK cells in the CXCR3^KO^ chimeras (**Figure S7A**). The resulting CXCR3^KO^- and CXCR3^WT^-chimeric mice were infected with LCMV prior to assessment of NK-cell localization on day 3 of infection (**Figure 3B**). Similar to infected non-chimeric C57BL/6 mice (WT), a substantial fraction of NK cells co-localized within the splenic white pulp (**Figure 3C**) and T-cell zones (**Figure 3D**) of infected CXCR3^WT^ BMC mice. In contrast, NK-cell frequencies within the splenic white pulp (**Figure 3C**) and T-cell zones (**Figure 3D**) were significantly reduced in CXCR3^KO^ BMCs. Thus, localization of NK cells within T-cell rich regions of the splenic white pulp during LCMV infection depends on NK-cell expression of CXCR3.

### CXCR3 is required for NK-cell suppression of anti-viral T cells

Transient localization of NK cells in the splenic white pulp during LCMV infection (**Figure 1**) coincides with the window of time during which NK cells kill activated T cells (5). Since CXCR3 is necessary for NK-cell localization within T-cell rich white pulp during LCMV infection (**Figure 3**), we hypothesized that mice lacking CXCR3 on NK cells would display an enhanced virus-specific T-cell responses similar to those in mice depleted of NK cells. Anti-NK1.1 antibody treatment one day before infection effectively depleted NK cells from both sets of mixed BMC mice (**Figure S7B**). At day 7 of LCMV infection, antiviral T-cell responses were assessed by intracellular cytokine staining after in vitro re-stimulation with viral peptide. The proportion (**Figure 4A**) and number (**Figure 4B**) of IFN-*γ*_+_TNF_+_ LCMV GP_64-80_-specific CD4 T cells was elevated to a similar extent in CXCR3^KO^ chimeras and in NK-cell depleted CXCR3^WT^ chimeras, with no apparent additive effect of NK cell-depletion in CXCR3^KO^ chimeras. The numbers of splenic Fas^+^GL-7^+^ germinal center B cells were similarly increased by CXCR3-deficiency or NK-cell depletion (**Figure 4C**).

**Figure 4.**
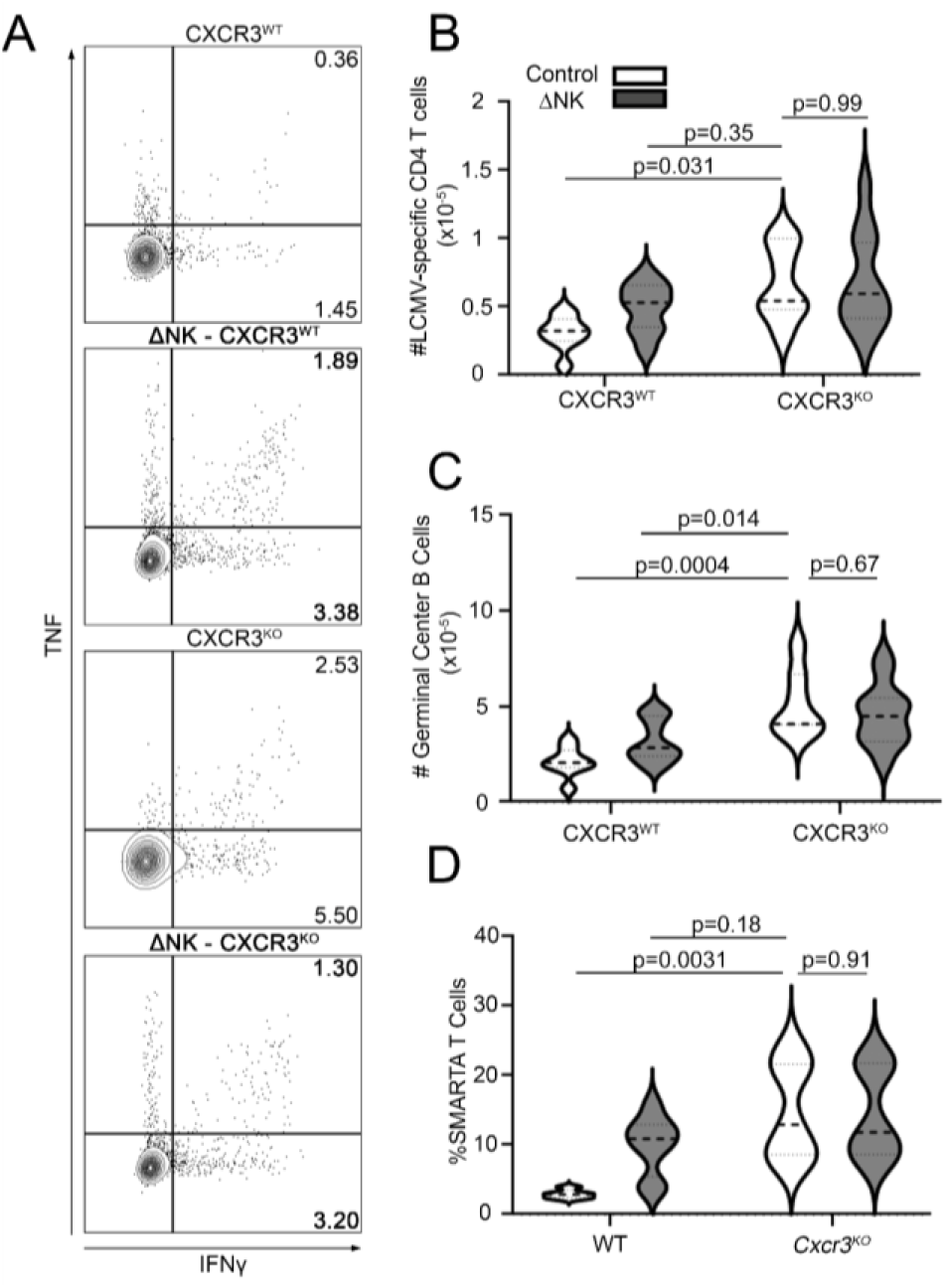
CXCR3 is required for NK-cell suppression of antiviral T cells. (A-C) Mixed CXCR3^WT^ and CXCR3^KO^ bone-marrow chimeric mice (n=7-12 mice/group) were depleted of NK cells using anti-NK1.1 antibody (ΔNK) or treated with control antibody one day prior to infection with LCMV. (B) Numbers of IFN-*γ*^+^ TNF^+^ expressing LCMV GP_64-80_-specific CD4 T cells were measured on day 7 by intracellular cytokine staining and flow cytometry. (C) Numbers of Fas^+^ GL-7^+^ germinal center B cells were also measured. (D) C57BL/6 and *Cxcr3*^*KO*^ (n=3-4 mice/group) mice were depleted of NK cells (ΔNK) or given non-depleting control antibody prior to intravenous infusion of 5×10^5^ LCMV-specific Ly5.1^+^ transgenic SMARTA CD4 T cells and infected one day later with LCMV. At day 6, the proportions of donor TCR-V*α*2^+^Ly5.1^+^CD4^+^ SMARTA T cells that were IFN-γ^+^ TNF^+^ were quantified in the spleens of recipient mice. Statistical analyses were performed using a one-way ANOVA. Data are a combination of two independent experiments.

As an alternative test of our hypothesis, CXCR3-sufficient LCMV-specific TCR transgenic SMARTA CD4 T cells (Ly5.1) were seeded in either CXCR3-sufficient C57BL/6 mice or germline *Cxcr3*^*KO*^ mice (Ly5.2) prior to NK-cell depletion and LCMV inoculation. On day 6 of infection, the frequencies of donor SMARTA cells expressing IFN-γ and TNF were increased in *Cxcr3*^*KO*^ and NK-cell-depleted recipient mice relative to those in control animals (**Figure 4D**). There was no additional effect of NK-cell depletion in *Cxcr3*^*KO*^ hosts. Thus, CXCR3 expression is important for NK-cell suppression of antiviral T cells.

## Discussion

These data reveal a crucial spatiotemporal mechanism of NK-cell regulation of antiviral T-cells. We identify CXCR3 as essential for both transient localization of NK cells within T-cell zones after infection and in the resulting suppression of antiviral T-cells by NK cells. We demonstrate that IFN-I-dependent induction of CXCR3 ligand expression is vital mechanism determining NK-cell localization in T-cell zones. Adenoviral vectors that do not robustly induce IFN-I also fail to elicit NK-cell follicular migration and associated suppression of T cells. Therefore, CXCR3 directs crucial redistribution of NK cells during inflammation that brings these cells into close proximity of activated T cells to permit perforin-dependent regulation (**Figure S8**). Translational targeting of this mechanism represents an innovative means of potentially enhancing vaccine efficacy and boosting antibody generation.

The transient nature of white pulp localization of NK cells represents a mechanistic explanation for the narrow temporal window of immunoregulatory function of NK cells. Measurable killing of activated T cells by NK cells is limited to the first three days of acute LCMV infection (4, 5), overlapping with the peak localization of NK cells in the white pulp reported here. Rapid egress or loss of NK cells from T-cell zones after day 3 likely limits further immunoregulatory killing. The degree to which different pathogens or vaccines trigger CXCR3-dependent migration of NK cells is linked to the ability to induce IFN-I responses. Some adenoviral vectors (e.g. Ad28 and Ad35) do trigger IFN-I release (32, 33), but these responses are several orders of magnitude lower than those seen after LCMV infection (34) and rapidly wane within hours of inoculation.

In addition to LCMV, re-localization of NK cells to T-cell zones (16, 18, 35) and NK-cell suppression of T cells (5, 36) are also triggered during infections with MCMV or after injection of TLR ligands. Induction of IFN and ligands for CXCR3 (37, 38) are shared features of these contexts. This explains why viruses that poorly stimulate NK-cell inhibition of T cells, including vaccinia virus (39) and Ad5 vectors (25), also weakly induce IFN responses (33, 40). Of note, transient blockade of IFN-I early during LCMV infection can enhance resulting antiviral T- and B-cell responses (41). Ablation of the IFNAR on NK cells exerts a similar effect (42). In MCMV and SIV/HIV infections, where NK cells exhibit potent antiviral function, this migration of NK cells to T-cell zones serves to protect these tissues from virus (17, 22). However, conservation of this migration during immunization or during infection with viruses that are refractory to NK-cell antiviral functions (e.g. LCMV) is more consequential for immune regulation than for immune defense.

Our discovery of CXCR3-mediated, white-pulp localizing NK cells in suppression of T cells opens new avenues of research that will enhance our understanding of the biology of this process as well as provide translational targets to circumvent this activity of NK cells. Immunosuppressive function of NK cells is linked to reduced T-cell memory (4), neutralizing antibody titers (4, 9), antibody affinity maturation (8), and vaccine efficacy (11). Our discovery highlights the potential for interventions blocking CXCR3 or its ligands in NK-cell immunosuppression to enhance vaccine efficacy. Further characterization of white-pulp localized NK cells is likely to reveal key mediators involved in T-cell suppression. Indeed, white-pulp and red-pulp localized NK cells exhibit markedly different transcriptomes (unpublished observations). This represents a promising pipeline for identification of translational targets to enhance vaccine-elicited immune responses by circumventing NK-cell activity.

## Materials and Methods

### Study approval

Experiments were performed according to ethical guidelines approved by the Institutional Animal Care and Use Committee and the Institutional Biosafety Committee of Cincinnati Children’s Hospital Medical Center. Additional methods details are presented in the Supplemental Material.

### Mice

C57BL/6, *Cxcr3*^KO^, B6.SJL-*Ptprc*^*a*^ *Pepc*^*b*^/BoyJ (Ly5.1), and B6.129S7-*Rag1*^*tm1*.*1Cgn*^/J (*Rag*^KO^) mice were purchased from Jackson Laboratory (Bar Harbor, ME). SMARTA LCMV-specific TCR transgenic mice were obtained from Shane Crotty and bred to Ly5.1 in house. Male mice between 8-20 weeks of age were routinely utilized in experiments. Mice were housed under barrier conditions. Staff were blinded to genotype and treatment status of experimental groups during harvesting, processing, and data acquisition.

### Infections and vector injections

Mice were infected with the Armstrong strain of LCMV via intraperitoneal (i.p.) injection of 5×10^4^ plaque forming units (p.f.u.) per mouse. Other mice were injected i.p. with 3×10^7^ p.f.u. adenovirus serotype 5 (Ad5)-LacZ or 1×10^10^ viral particles of Ad5-LCMV GP.

### Mixed bone marrow chimeric mice

Mixed bone marrow chimeric (BMC) mice deficient in or expressing CXCR3 were generated by admixing the donor bone marrow of *Rag*^*KO*^*Cxcr3*^*WT*^ (referred to as CXCR3-WT) or *Rag*^*KO*^*Cxcr3*^*KO*^ (referred to as CXCR3-KO) mice at a 9:1 ratio with bone marrow from wild-type Ly5.1 mice (31). In this mixture, NK cells and other innate lymphocytes preferentially derive from *Rag*^*KO*^ precursors while the adaptive compartment (T and B cells) can only develop from Ly5.1 precursors and as such are always CXCR3^WT^. Bilateral femur bones of mice were harvested and stripped of muscle tissue using gauze sponges and blades. Femur heads were cut to allow thorough flushing of the bone marrow with 1X PBS into a petri dish using a 10mL syringe topped with a 26G needle. Bone marrow cells were counted using a hemocytometer and a bone marrow cell suspension was prepared to achieve 10×10^6^ cells/mouse. Recipient C57BL/6 mice were lethally irradiated (See Irradiation) prior to intravenous injection of mixed bone marrow cells. A minimum of 16 weeks was allotted for reconstitution prior to initiating experimental infections.

### SMARTA transfer model

SMARTA mouse spleens were harvested and processed into single cell suspensions (See Flow Cytometry) from which CD4 T cells were enriched by depletion of non-CD4 T cells (Miltenyi). 5×10^5^ CD4 T cells were retro-orbital (r.o.) injected into CXCR3-WT, CXCR3-KO and NK-depleted mice 1 day prior to LCMV infection. Six days following infection, spleens were harvested and SMARTA^+^ CD4 T cell numbers, activation status and cytokine production were assessed by flow cytometry following *in vitro* LCMV-GP_64-80_ (GPDIYKGVYQFKSVEFD) peptide stimulation.

### Irradiation

Irradiation of mice was performed using the J. L. Shepherd Mark I Cesium Irradiator. Gamma irradiation was provided by Cesium 137 Source. Mice were placed into a rotating pie-shaped holder to limit mobility and ensure equal irradiation. The pie-shaped holder was then placed in the irradiator to deliver the required dose at a dose rate of 0.5 Gy/min. The optimal dose for hematopoietic ablation C57BL/6 mice is 11.75 Gy. All lethal doses/ablative doses were delivered as two split exposures to limit non-hematopoietic toxicity. The split exposures of 7.0 Gy for first exposure and 4.75 Gy for the second were separated by 3 hours. No anesthesia or pain meds were used in this procedure.

### Intravascular staining

Protocols for intravascular staining of lymphocytes (23) were adapted to permit staining of NK cells. A few minutes prior to harvest, mice were anesthetized with isoflurane then given (r.o.) injections of 1.2 μg of anti-NKp46-APC (29A1.4, Biolegend) or goat anti-Mouse anti-NKp46 (R&D, AF2225) in 200 μL PBS or HBSS per mouse. The injected antibody was allowed to circulate in the mice for no more than 3 minutes, after which mice were euthanized. Spleens were then harvested, processed, and stained as described under flow cytometry or prepared for confocal imaging as described under tissue processing, sectioning and immunohistochemistry.

### In vivo NK-cell depletion, recombinant IFN-α and IFNAR-1 blockade

One or two days before infection, selective depletion of NK cells was achieved through a single i.p. injection of 25 μg anti-NK1.1 monoclonal antibody (PK136) or 25 μg of a control mouse IgG2a isotype antibody (C1.18.4) produced by Bio-X-Cell (West Lebanon, NH). At time of Ad5-GP or -LacZ vector immunization and the two consecutive days following, i.p. injection of 8 μg/mouse of recombinant interferon-α from Biolegend (752806) was administered. Similarly, on days 0, 1, and 2 of LCMV infection, IFN-I signaling was blocked via i.p. injection of 400 μg/mouse anti-IFNAR-1 (MAR1-5A3) antibody from Bio-X-Cell (West Lebanon, NH). In other experiments, mice were given one i.p. injection of 100 μg/mouse anti-IFNAR-1 antibodies alongside LCMV infection.

### RNA Isolation, cDNA synthesis and RT-qPCR

Spleens harvested from mice were processed into single cell suspensions as described below (Flow Cytometry). 30mg of whole spleen RNA was extract via 30 second homogenization in RLT lysis buffer containing 1% β-mercaptoethanol per RNeasy kit (Qiagen). RNA was reverse transcribed into cDNA using iScript cDNA synthesis kit (BioRad). Subsequent RT-qPCR was performed using Taqman Gene Expression Assays to detect *Cxcl9* (Mm00434946_m1), *Cxcl10* (Mm00445235), *Cxcl11* (Mm00444662_m1), *Ifna9* (Mm00833983_s1), *Ifnb1* (Mm00439552_s1) and *Gapdh* (Mm99999915_g1), and Taqman Fast Advance master mix (ThermoFisher), run using Applied Biosystems 7500 Real-Time PCR (ThermoFisher) and 7500 software v2.3.

### Flow cytometry and in vitro peptide stimulation

Single-cell leukocyte suspensions were prepared from spleens by mechanical homogenization of tissues between frosted glass microscope slides (VWR) and filtration through a 70µm nylon mesh. Following lysis of red blood cells, cells were plated at 2×10^6^/well in 96-well round-bottom plates, then either immediately subjected to flow staining or first stimulated for 5 hours at 37°C with 1 μM LCMV-GP_64-80_ peptide in the presence of brefeldin A. Leukocytes were then washed and resuspended in FACS buffer (HBSS or PBS + 2-5% fetal bovine serum + 0.5 mM EDTA) prior to surface staining with the following antibodies: CD3e (145-2C11), CD19 (1D3), TCRβ (H57-597), NKp46 (29A1.4), NK1.1 (PK136), CD49b (DX5), CD4 (GK1.5), CD8α (53-6.7), CD44 (IM7), CD43 (1B11), TCRVα2 (B20.1), CD45.1 (A20), CD45.2 (104), CXCR3 (CXCR3-173), Fas (SA367H8) IgD (11-26c.2a), GL-7 (GL7), NKG2D (1D11), KLRG1 (2F1/KRLG1), NKG2A (16A11), CD94 (18d3), Ly49H (3D10), DNAM-1 (TX42.1), CD11b (M1/70), CD27 (LG.3A10), CD25 (PC61) and granzyme B (GB11). All antibodies were purchased from Biolegend or BD Biosciences. Cells were surface stained for 15-30 minutes at 4°C. For CXCR3 staining, cells were incubated at room temperature. Following staining, cells were washed and fixed with BD fixation buffer (BD Biosciences) for 5 minutes at 4°C. Cells were washed twice and resuspended in 200 μL FACS buffer. For intracellular staining, cells were washed and fixed with BD fixation & permeabilization buffer (BD Biosciences) for 15-20 minutes at 4°C. Cells were then intracellularly stained with anti-IFN-γ (XMG1.2) and anti-TNF-α (MP6-XT22) for 15-30 minutes at 4°C. Cells were then washed and resuspended in 200 μL FACS buffer and analyzed on a BD LSR Fortessa cytometer.

### Tissue processing, sectioning, and immunohistochemistry

Tissues to be analyzed by fluorescence microscopy were placed into 4% formaldehyde solution (EMS) for 5 hours followed by overnight dehydration in a 30% sucrose (Sigma) solution at 4°C. Samples were washed with 1X PBS prior to embedding within optimal cutting temperature (OCT) media (Sakura) and frozen using a dry ice slurry in 100% ethanol and stored in −20°C. Tissues were sectioned (7-10µm thick) using a cryostat (Leica CM3050 S) and affixed to positively charged slides (Denville). Slides were dried at RT for 5-10 minutes preceding a 10-minute fixation in chilled 100% acetone at −20°C. Slides were subsequently dried at RT for 5-10 minutes and washed twice in 1X chilled PBS before being placed in saturation buffer containing 10% normal donkey serum (Sigma), 0.1% Triton-X (Sigma) and 1X PBS for blocking at RT for 45 minutes. Slides were incubated with a primary antibody cocktail containing one or more of the following antibodies overnight at 4°C: goat anti-mouse NKp46 (1:100, R&D Systems), anti-mouse CD3e-AF594 or AF647 (500A2, 1:200) and anti-mouse CD169-AF594 or AF647 (3D6.112). The following day, slides were washed twice in 1X chilled PBS. Subsequent secondary antibody staining with donkey anti-goat AF488 (Abcam, 1:1000) was used for 2 hours at room temperature to reveal NKp46 primary staining. Slides were again washed twice with 1X chilled PBS, mounted with prolong diamond mounting media (Thermo-Scientific), covered with a coverslip (Thermo), and allowed to cure overnight prior to imaging.

### Confocal Microscopy & NK-cell quantification

Confocal imaging was performed using a Laser Scanning Nikon A1RSi Inverted Confocal Microscope with NIS Elements Confocal software. Z-stacked tissue images were acquired through a 10X objective (Nikon Plan Apo λ) from which a maximum intensity projection is generated prior to cell enumeration. Using tools available in NIS Elements Analysis software and guided by CD169- and CD3-staining, borders around white pulp (CD169 boundary) and T cell zones (CD3 boundary) were drawn. B220- and CD3-staining were used to delineate borders when enumerating NK cells within lymph nodes. NK cells located with these borders were enumerated using unbiased NIS Elements-derived algorithms and plotted using GraphPad Prism (San Diego). To calculate NK cell proportions within the white pulp or T-cell zone, the total NKp46^+^ cells within the respective compartment were normalized to total NKp46^+^ cells within the field of view. Brightness and contrast for each representative image were adjusted equally across all channels using Photoshop CS6.

#### Statistical Analysis

Experimental results are consistently presented as the median with individual data point spread. Statistical differences between control and experimental groups were calculated using a one-way analysis of variance (ANOVA) with either the Holm-Šídák multiple comparison test or a Dunnett’s multiple comparisons test. When a confidence interval was not desired, the Holm-Šídák multiple comparisons method was used to generate higher statistical power. A p-value of less than 0.05 was considered significant. Graphing and statistical analysis were routinely performed using GraphPad Prism (San Diego). Researchers were blinded to experimental groups during

## Acknowledgements

We thank Dr. Kofron and the Comprehensive Mouse and Cancer Core for support, Dr. Molkentin for Ad-LacZ, Dr. Crotty for SMARTA mice, Shane D’Souza for crafting of the summary model, and Drs. Kottyan, Lewkowich, Hildeman, Herro, and Borchers for critical reading of the manuscript. Funding for this work was provided NIH grants DA038017, AI148080, AR073228 (SNW), T32GM063483 (AA, NL, HAF), and T32AI118697 (AA). Support was also provided by the Cincinnati Children’s Research Foundation (SNW), L.B. Research and Education Foundation (N.L.), and Albert J. Ryan Foundation (AA). The Flow Cytometry Core was supported by NIH AR070549 and P30 DK078392.

---

**Supplemental Figure 1.**
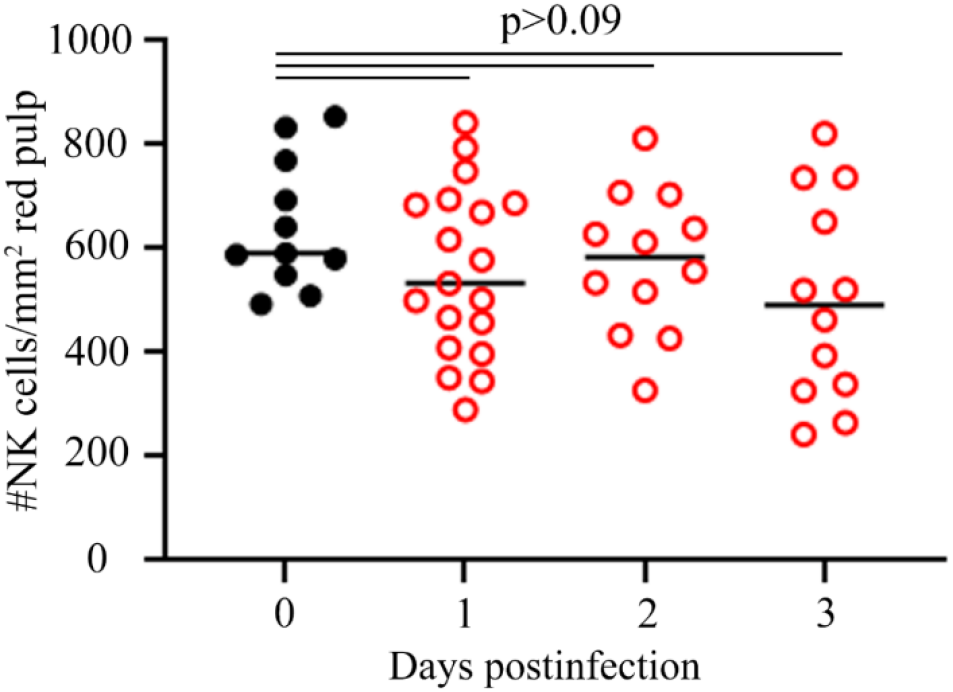
Enumeration of NKp46^+^ NK cells within the splenic red pulp following infection. C57BL/6 mice (n=3-4/group) were LCMV-infected. At the indicated time-point post infection, the number (5-12 follicles/mouse) of NKp46^+^ NK cells (green) were enumerated within the splenic red pulp defined by the region not bound the CD169 (purple) macrophages. Data are representative of two independent experiments.

**Supplemental Figure 2.**
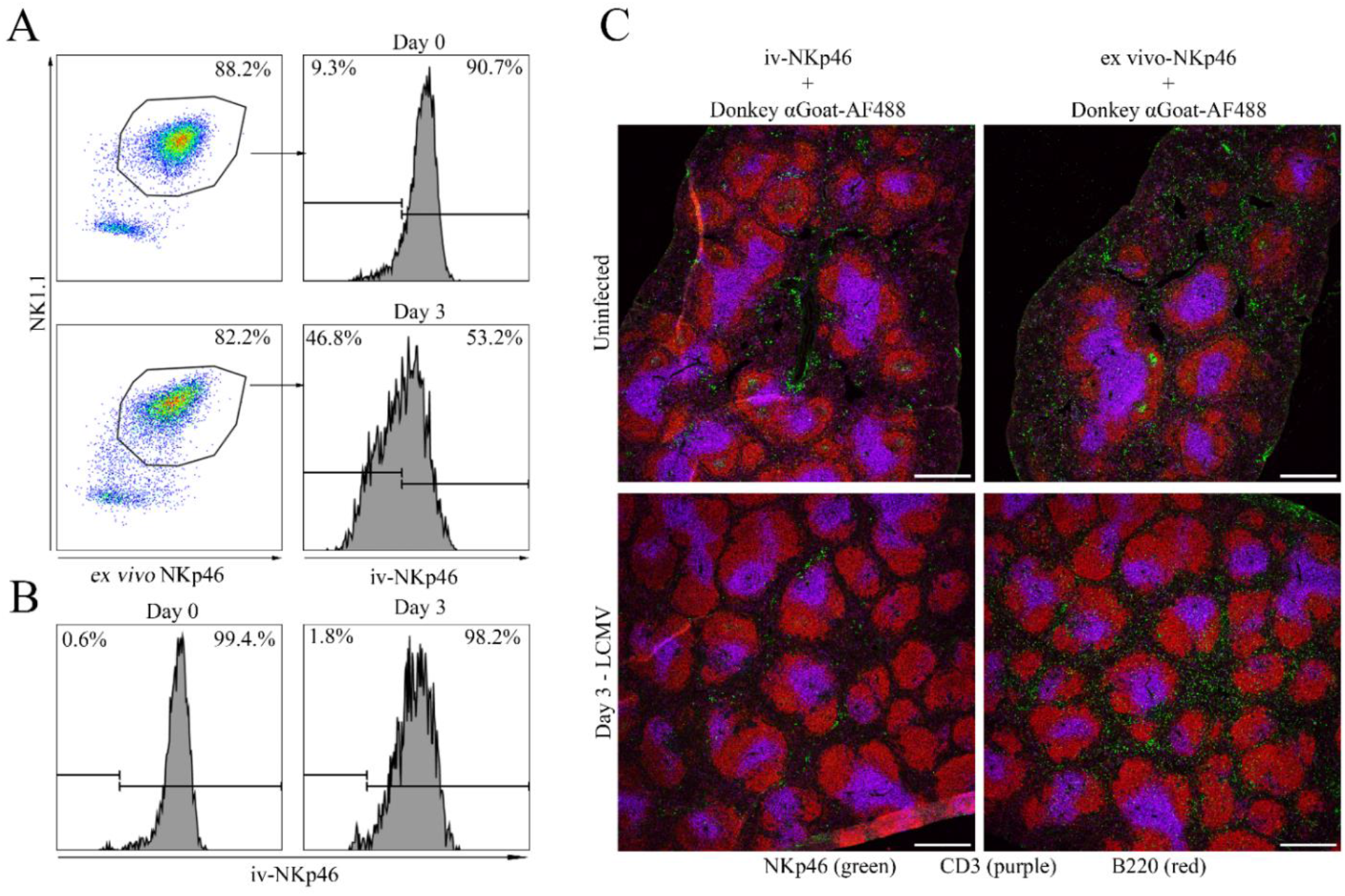
Ex vivo and in vivo NKp46 staining in blood and spleen NK cells. (A) Representative histograms showing iv-NKp46 expression of circulating (CD3^neg^ TCRβ^neg^ CD8α^neg^ CD49b^+^ NK1.1^+^ ex-vivoNKp46^+^) NK cells at indicated times post LCMV infection. (B) Representative plots show iv-NKp46 staining on NK cells (CD3^neg^ TCRβ^neg^ CD8α^neg^ CD49b^+^ NK1.1^+^ ex-vivoNKp46^+^). (C) Representative confocal images of spleen sections from uninfected mice and infected mice 3 days following LCMV injection visualizing NKp46^+^ (green) NK cells, B220^+^ (red) B cells and CD3^+^ (purple) T cells following staining with either donkey anti-goat-AF488 secondary antibody alone or re-stained with goat anti-mouse anti-NKp46 antibody ex vivo prior to donkey anti-goat secondary. Scale bars measure 500 µm.

**Supplemental Figure 3.**
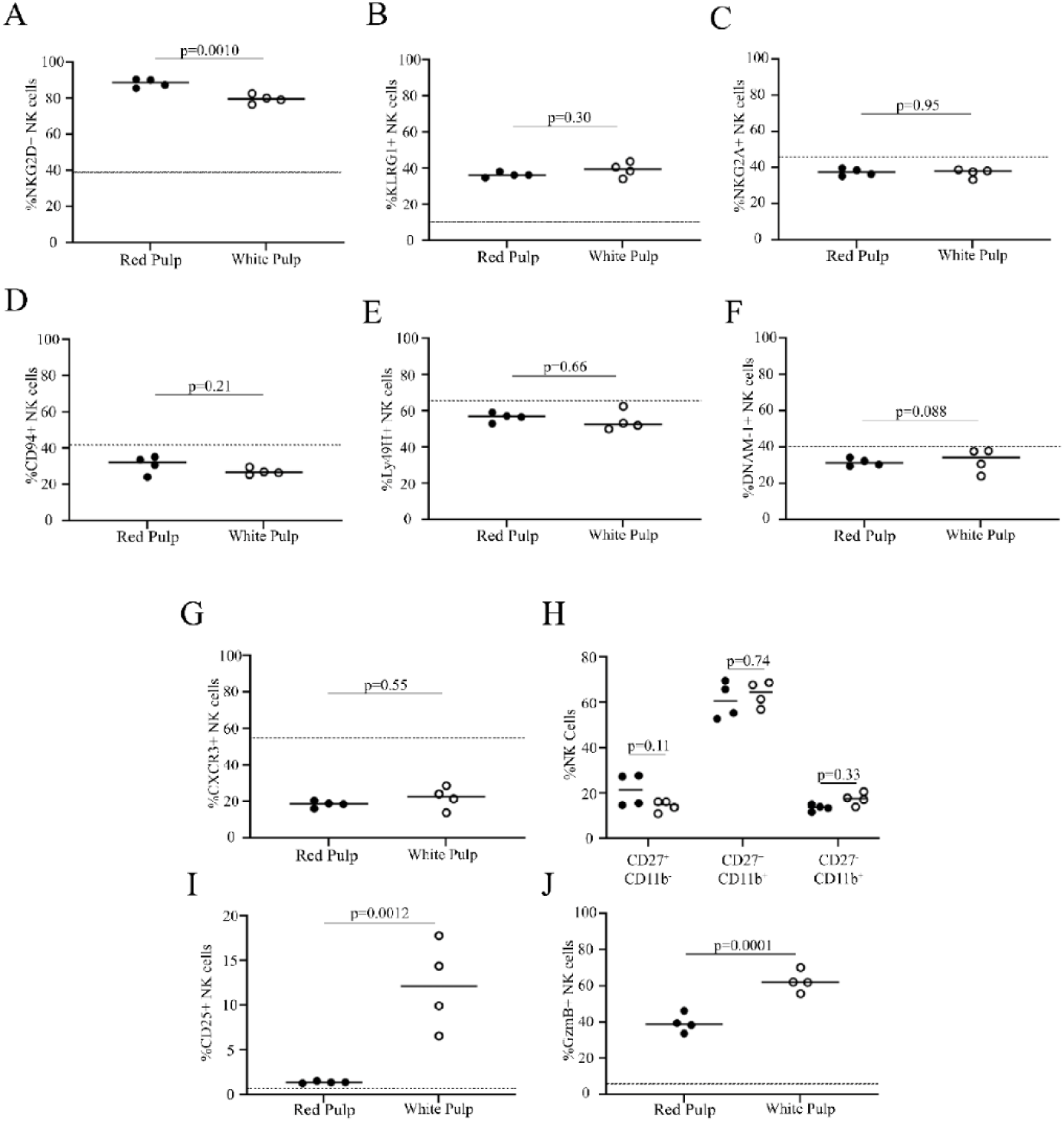
Characterization of red pulp and white pulp-localized NK cells. (A-K) Three days following LCMV infection, C57BL/6 (n=4) mice were intravenously injected with anti-NKp46 antibody three minutes prior to euthanasia to label splenic NK cells (CD3^neg^ TCRβ^neg^ CD8α^neg^ CD49b^+^ NK1.1^+^ ex-vivoNKp46^+^) in vascularized red pulp (iv-NKp46^+^) or white pulp (iv-NKp46^neg^) regions. Cells were stained for the indicated markers and assessed by flow cytometry. The proportion of white pulp or red pulp NK cells expressing the indicated markers is shown. Dotted line reflects expression level in uninfected mice. Data reflect one of two independent experiments with statistical differences determined by one-way ANOVA.

**Supplemental Figure 4.**
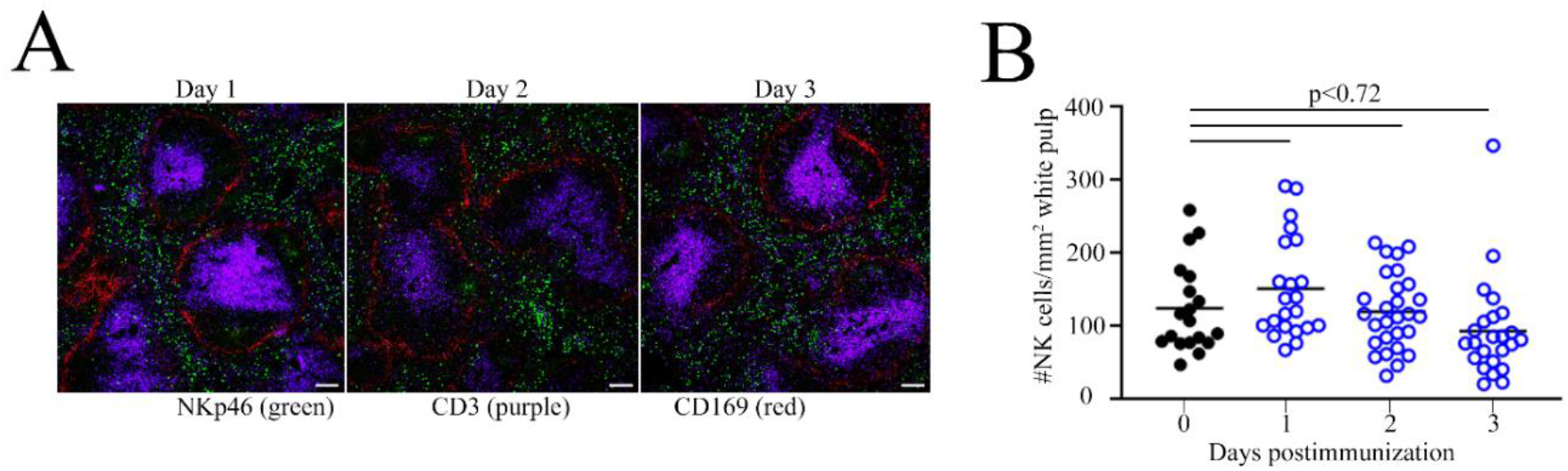
Splenic NK cell migration during adenovirus vector immunization. (A-B) C57BL/6 mice were inoculated with Ad5-LaZ. At indicated times post inoculation, spleens were imaged to determine localization of NKp46^+^ NK cells (green) relative to CD169^+^ macrophages (red), with enumeration of NK cells therein. Scale bars measure 100 µm.

**Supplemental Figure 5.**
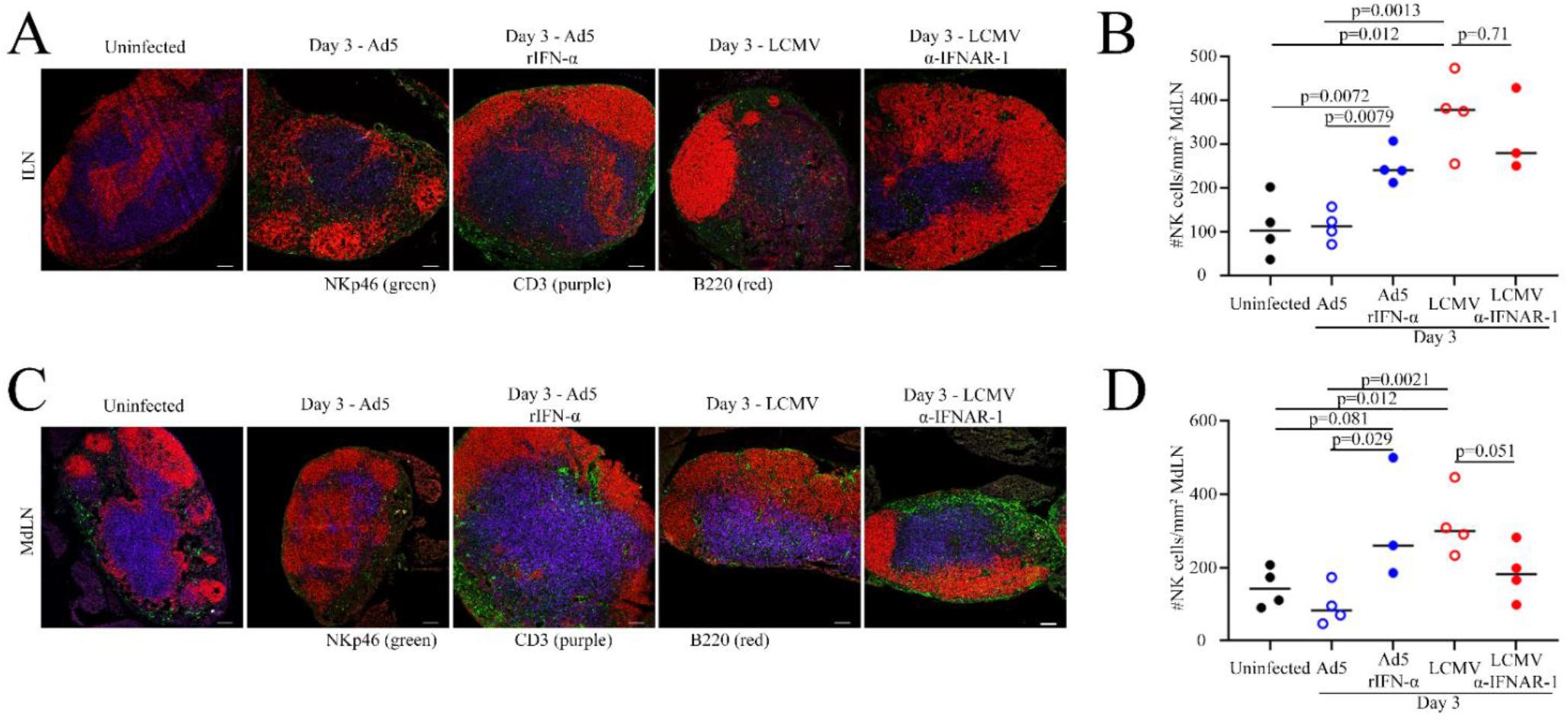
Role of type I interferons in NK cell localization in draining lymph nodes. (A-D) C57BL/6 mice (n=3-4/group) were inoculated with Ad5-LacZ, LCMV or uninfected and treated with either 8 μg/mouse recombinant IFN-α or 400 μg/mouse anti-IFNAR-1 antibodies. NKp46^+^ (green) NK cells were enumerated within the B220^+^ (red) and CD3^+^ inguinal (A-B) and mediastinal (C-D) lymph nodes at the indicated time point. Data reflect one of two independent experiments with statistically significant differences determined by one-way ANOVA. Scale bars measure 100 µm.

**Supplemental Figure 6.**
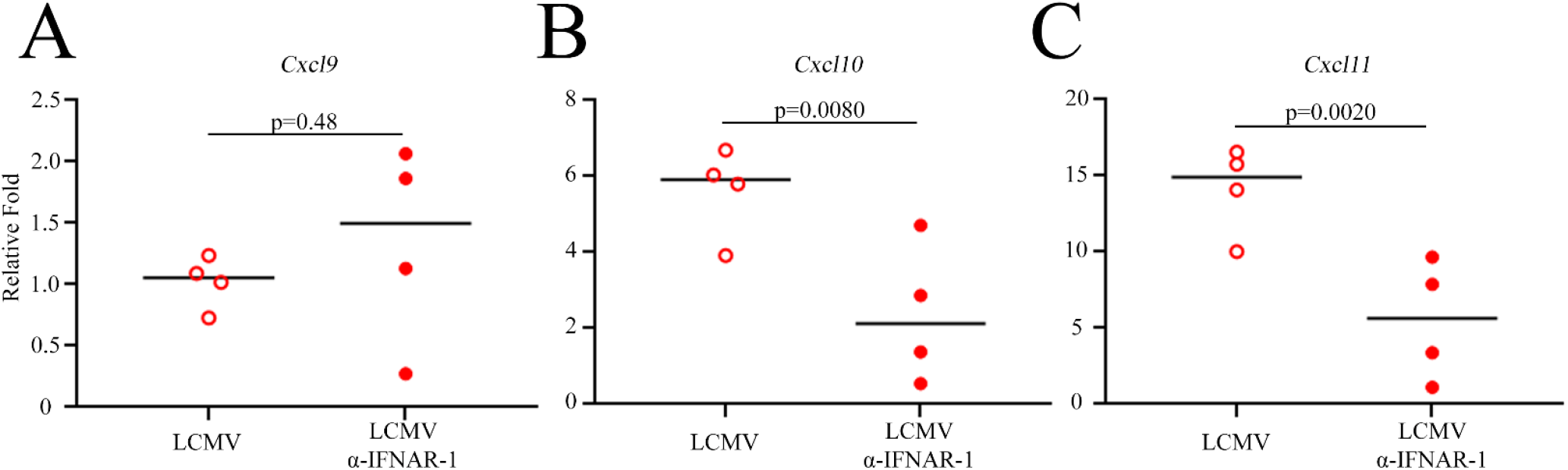
Reduced CXCR3 ligand expression following IFNAR blockade during LCMV infection. C57BL/6 mice (n=4/group) were treated or not with 100 μg anti-IFNAR antibodies prior to LCMV infection. Two days following infection, spleens were harvested, and mRNA was extracted for RT-qPCR analysis of *Cxcl9, Cxcl10*, and *Cxcl11* expression levels. Statistical differences determined by Student’s T-test.

**Supplemental Figure 7.**
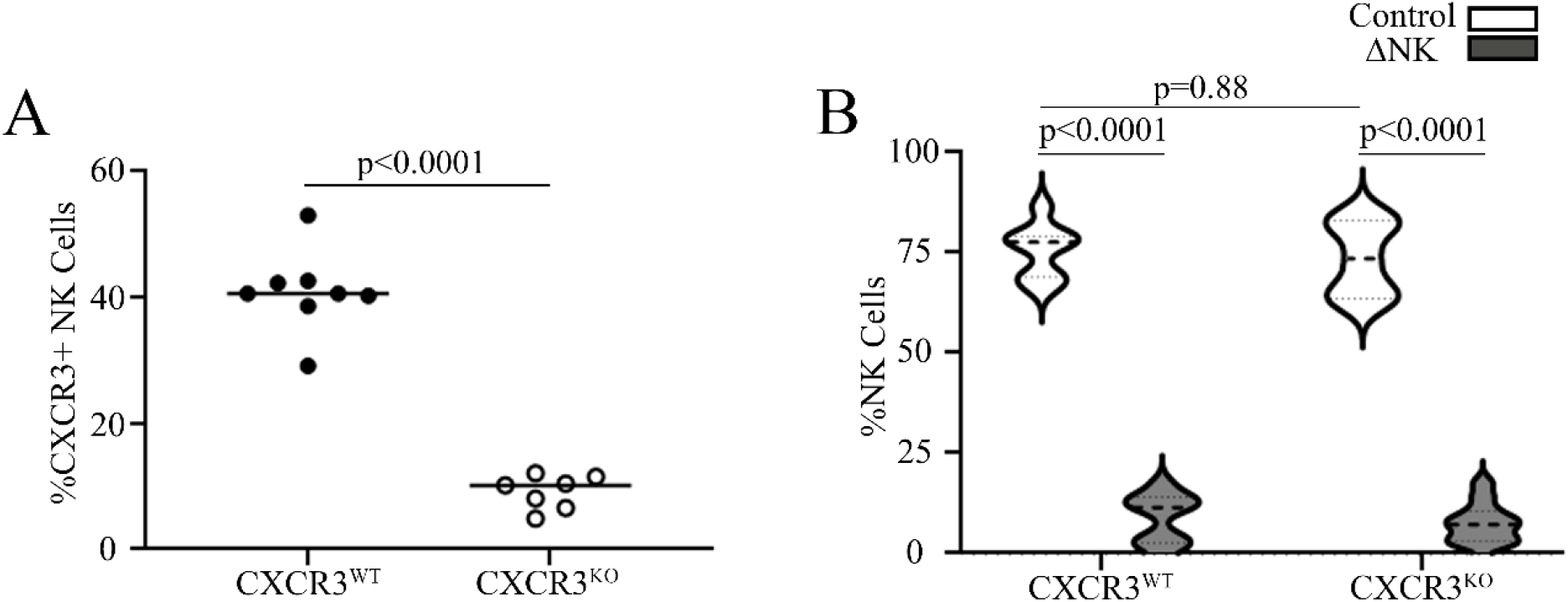
Confirmation of NK cell depletion and CXCR3 deletion in NK cells. Proportion of CD49b^+^ NKp46^+^ NK1.1^+^ NK cells in mixed CXCR3^WT^ or CXCR3^KO^ bone-marrow chimeras that (A) express CXCR3 or were (B) depleted or not of NK cells. Data reflect a combination of two independent experiments with statistically significant differences determined by one-way ANOVA or Student’s T-test.

**Supplemental Figure 8.**
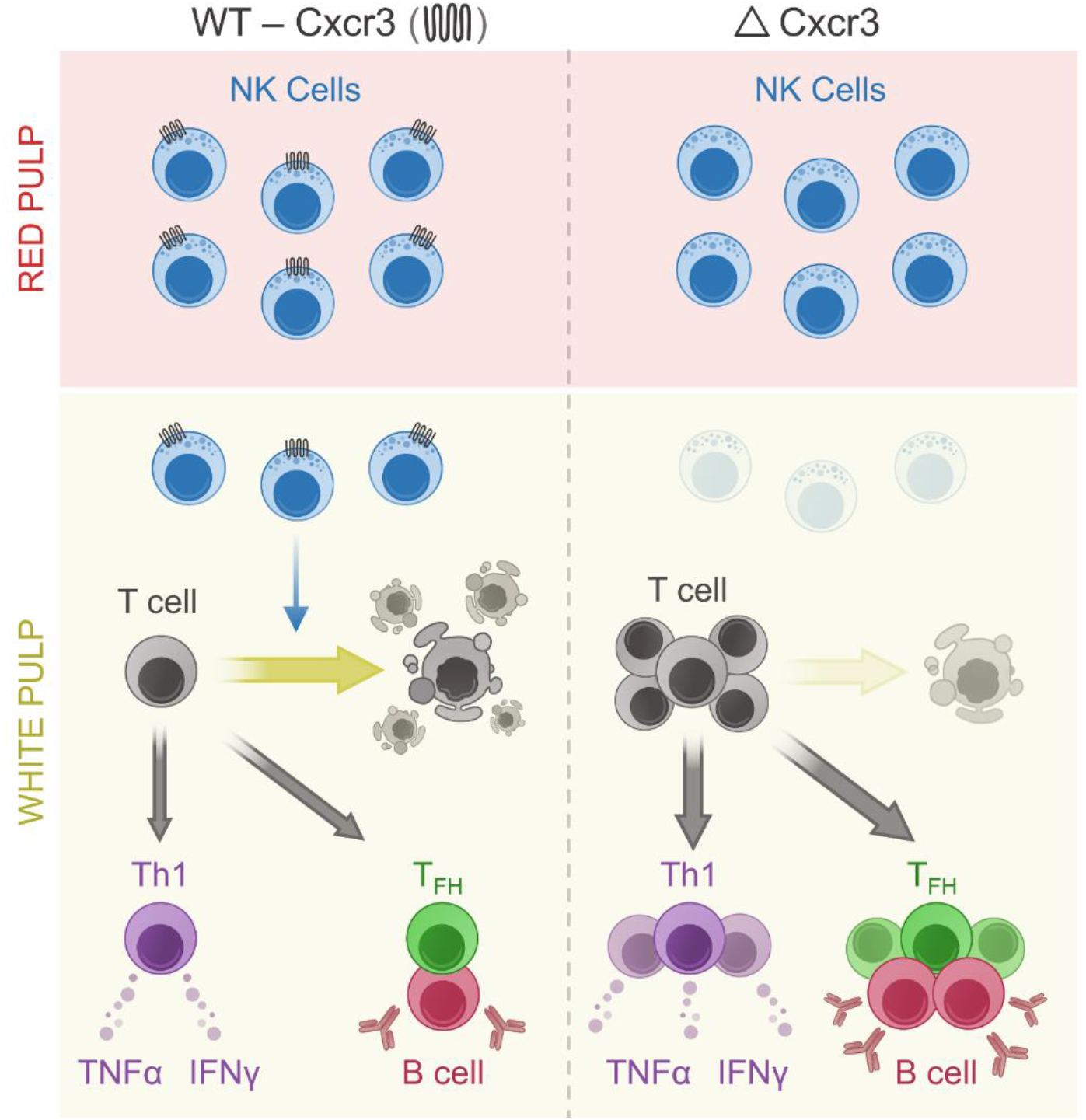
Model of CXCR3-dependent NK cell immunoregulatory activity during infection. Viral infection leads to NK cell localization within T cell-rich follicles wherein they cull antiviral T cells, diminishing T cell cytokine responses and generation of TFH cells. We propose that loss of CXCR3 on NK cells impairs this localization, rescuing T cells from NK cell-driven death thereby allowing for increased T cell cytokine responses and TFH cell generation important for viral control and antibody production.

## Notes

### Competing Interest Statement

The authors have declared no competing interest.

